# A Lentiviral Envelope Signal Sequence is a Tetherin Antagonizing Protein

**DOI:** 10.1101/2021.08.19.457005

**Authors:** James H. Morrison, Eric M. Poeschla

**Affiliations:** Division of Infectious Diseases, University of Colorado Anschutz Medical Campus, Aurora, CO 80045, USA

**Keywords:** Tetherin, BST2, HIV-1, FIV, restriction factor, accessory protein, innate immunity, Signal Peptide, Signal Sequence, Leader Sequence

## Abstract

Signal sequences are N-terminal peptides, generally less than 30 amino acids in length, that direct translocation of proteins into the endoplasmic reticulum and secretory pathway. The envelope glycoprotein (Env) of the nonprimate lentivirus Feline immunodeficiency virus (FIV) contains the longest signal sequence of all eukaryotic, prokaryotic and viral proteins (175 amino acids). The reason is unknown. Tetherin is a dual membrane-anchored host protein that inhibits the release of enveloped viruses from cells. Primate lentiviruses have evolved three antagonists: the small accessory proteins Vpu and Nef, and in the case of HIV-2, Env. Here we identify the FIV Env signal sequence (Fess) as the FIV tetherin antagonist. A short deletion in the central portion of Fess had no effect on viral replication in the absence of tetherin but severely impaired virion budding in its presence. Fess is necessary and sufficient, acting as an autonomous accessory protein with the rest of Env dispensable. In contrast to primate lentivirus tetherin antagonists, it functions by stringently blocking the incorporation of this restriction factor into viral particles rather than by degrading it or downregulating it from the plasma membrane.

## INTRODUCTION

The evolution of host antiviral factors has selected for reciprocal evolution of viral countermeasures, which can act through passive avoidance or direct antagonism. Tetherin (BST-2) is a type I interferon (IFN) inducible protein that forms homodimers and directly links newly formed HIV-1 particles and the plasma membrane through its transmembrane domain and a C-terminal GPI-anchor (Neil et al., 2008; Van Damme et al., 2008). This attachment function prevents viral particle release from infected cells and can lead to virus internalization, degradation via endosomal/lysosomal pathways, and induction of NFκB-dependent pro-inflammatory responses in the infected cell (Cocka and Bates, 2012; Galao et al., 2012; Miyakawa et al., 2009).

The importance of tetherin evasion for primate lentiviruses is indicated by their nearly ubiquitous encoding of antagonists. Three different such proteins have been described. SIVs of *Cercopithecus* genus primates (SIVgsn, SIVmus and SIVmon) and HIV-1 counteract tetherin with the accessory protein Vpu (Sauter et al., 2009). SIVcpz, the proximate precursor to HIV-1, shares common ancestry with cercopithecine SIVs yet utilizes Nef to counteract tetherin (Sauter et al., 2009). Other SIVs also utilize Nef to antagonize tetherin, including SIVsmm, the virus proximately ancestral to HIV-2 (Hirsch et al., 1989; Jia et al., 2009; Zhang et al., 2009). A small deletion in the cytoplasmic tail of human tetherin prevents Nef binding (Sauter et al., 2009). HIV-2 re-gained tetherin antagonism in its envelope glycoprotein (Le Tortorec and Neil, 2009), whereas the HIV-1 subgroups, which arose from independent cross-species transmission events, vary in this regard. Non-pandemic HIV-1 group O strains lack an efficient anti-tetherin mechanism, but pandemic HIV-1 group M strains evolved a Vpu capable of counteracting tetherin (Sauter et al., 2009). These primate lentiviral proteins all act by functionally depleting tetherin from the plasma membrane via intracellular sequestration or endocytosis and lysosomal degradation of the protein (Jia et al., 2009; Le Tortorec and Neil, 2009; Zhang et al., 2009).

For non-primate lentiviruses, much less is known about viral interaction with and evasion of tetherin. Their accessory gene repertoires are apparently more limited and Vpu and Nef are found only in primate lentiviruses. For that matter, no new lentiviral accessory genes have been identified for decades. Cat and dog tetherin proteins both restrict HIV-1 and FIV and both carnivore proteins are antagonized by FIV Env (Morrison et al., 2014). While this situation superficially resembles the antagonism of human tetherin by HIV-2 Env, major differences were observed that suggest different mechanisms. Unlike primate lentiviral antagonists, we found that the FIV Env mechanism does not require processing of Env into its surface unit (SU) and transmembrane (TM) domains (Morrison et al., 2014). It also shields the budding particle without downregulating plasma membrane tetherin, and does not rescue non-cognate (e.g., HIV-1) virus budding (Morrison et al., 2014). Here we explored the mechanism of FIV tetherin antagonism further and determined that it derives specifically from the signal sequence, which functions autonomously from Env, and acts to prevent particle incorporation.

## RESULTS

### Mapping of determinants of tetherin antagonism

We previously reported that the envelope glycoprotein (Env) of FIV counteracts restriction of this virus by both domestic cat and dog tetherin proteins and that this activity is independent of proteolytic processing of Env into surface unit (SU) and transmembrane (TM) domains (Morrison et al., 2014). FIV Rev and Env have the same initiator methionine codon but are differentiated by alternative splicing; therefore, Rev and Env of FIV share the first 80 amino acids (**Figure 1A**). To identify the minimal components of Env necessary to enable nascent FIV virion escape from tetherin-expressing cells, we constructed a series of Env frame-shift (efs) mutants that progressively truncate the protein while leaving Rev intact. These were constructed in FIVC36, an infectious molecular clone that replicates to high levels in vivo and causes feline AIDS (de Rozieres et al., 2004). Antagonism of tetherin was determined by quantifying FIV particles in cell supernatants following co-transfection of the FIV proviral construct and tetherin plasmids (**Figure 1B**). Co-transfection of an Env-intact FIV with increasing amounts of feline tetherin resulted in a modest reduction in reverse-transcriptase (RT) activity and capsid (CA) in supernatants (**Figure 1B**, blue bars and supernatant immunoblot). Introduction of a frameshift in the signal sequence of Env (amino acid 90) resulted in a virus that was significantly more sensitive to the presence of tetherin, with viral budding decreased in proportion to feline tetherin input (**Figure 1B**, orange bars and immunoblot). In contrast, termination of Env in mid-SU, at residue 330, reverted the phenotype, whereby again FIV was only modestly affected by tetherin co-expression (**Figure 1B**, grey bars).

**Figure 1.**
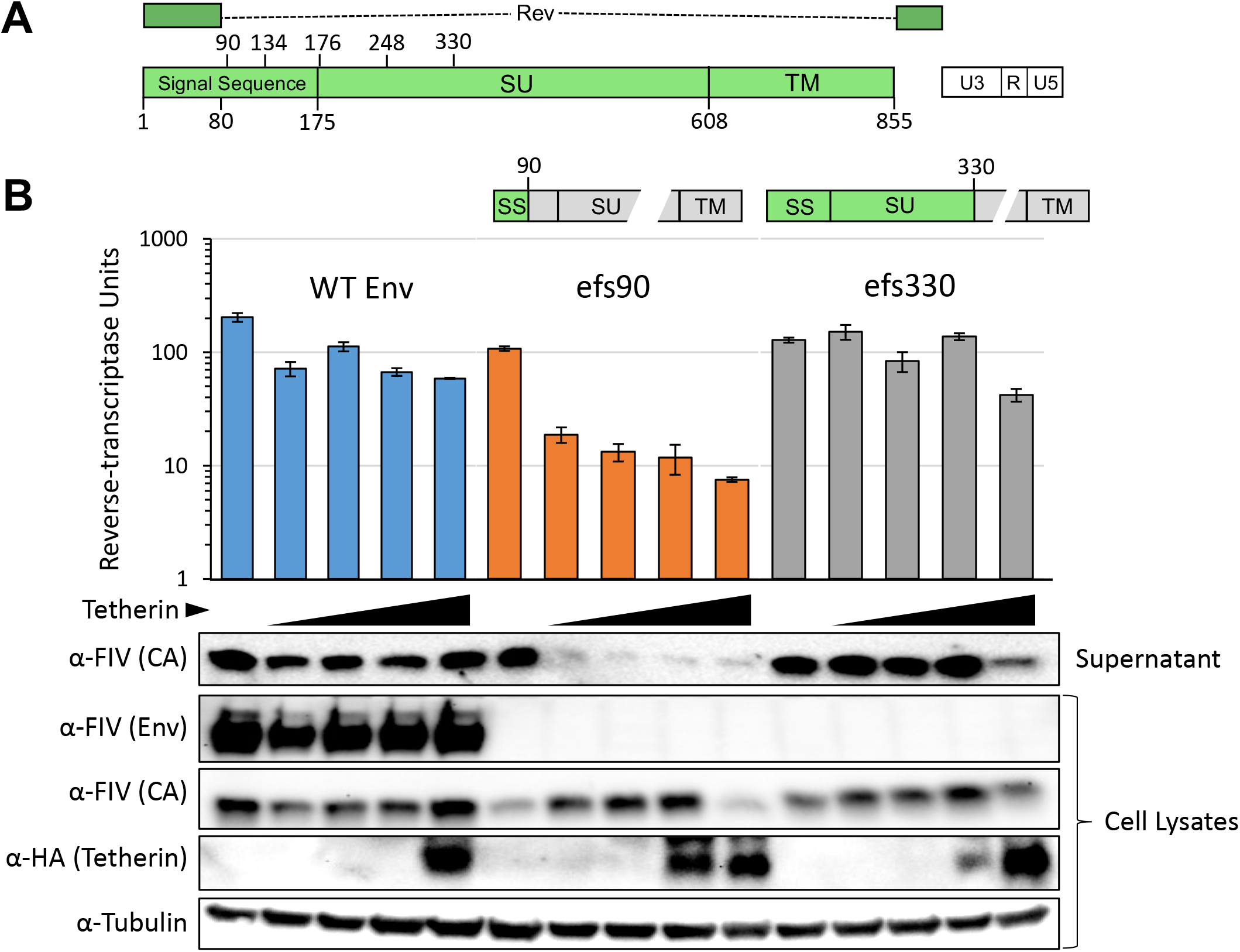
SU and TM are dispensable for tetherin antagonism. **(A)** Diagram of FIV *Env* gene. Numbering indicates amino acids from the Env initiator methionine, which is shared with Rev. **(B)** 293T cells were co-transfected with pFIVC36 proviral constructs (pCT-C36^A+^ based) encoding an intact Env (blue bars), or an Env frame-shift (efs) mutation at amino acid 90 (orange bars) or at amino acid 330 (grey bars). Diagrams indicate Env subunits intact (green) or disrupted (grey). Cell lysates and supernatant were harvested 48 hours after transfection and immunoblotted with the indicated primary antibody and a corresponding HRP-conjugated secondary antibody. Experiment was repeated four times and a representative example is shown.

To further map the virus-rescuing activity in Env, a set of three additional truncations was made by introducing frameshifts within the N-terminal portion of Env (efs134, efs176 and efs248). These plus the initial efs90 and efs330 viruses were tested for tetherin susceptibility, this time using cells that stably express either human or feline tetherin (**Figure 2A**). As expected, each viral variant expressed equivalent intracellular levels of FIV core proteins, budded in the absence of tetherin, and was blocked from budding by human tetherin (**Figure 2A**). Absence of the Env SU or TM domains did not significantly affect the ratio of intracellular Gag/CA to budded particles (efs176, efs248 and efs330). It was only when the signal sequence of Env was truncated (efs134 and efs90) that a loss of FIV budding was observed in cells that express domestic cat tetherin; again, human tetherin restriction was not abrogated) (**Figure 2A**). Quantification of immunoblot densities confirmed that when normalized for intracellular capsid expression levels feline tetherin was effective at blocking FIV budding of efs90 and efs134, whereas wild-type FIVC36 and the efs mutants retaining at least the first 176 amino acids of Env were resistant to feline tetherin and even had higher ratios of released to intracellular capsid compared to control cells lacking tetherin (**Figure 2B**).

**Figure 2.**
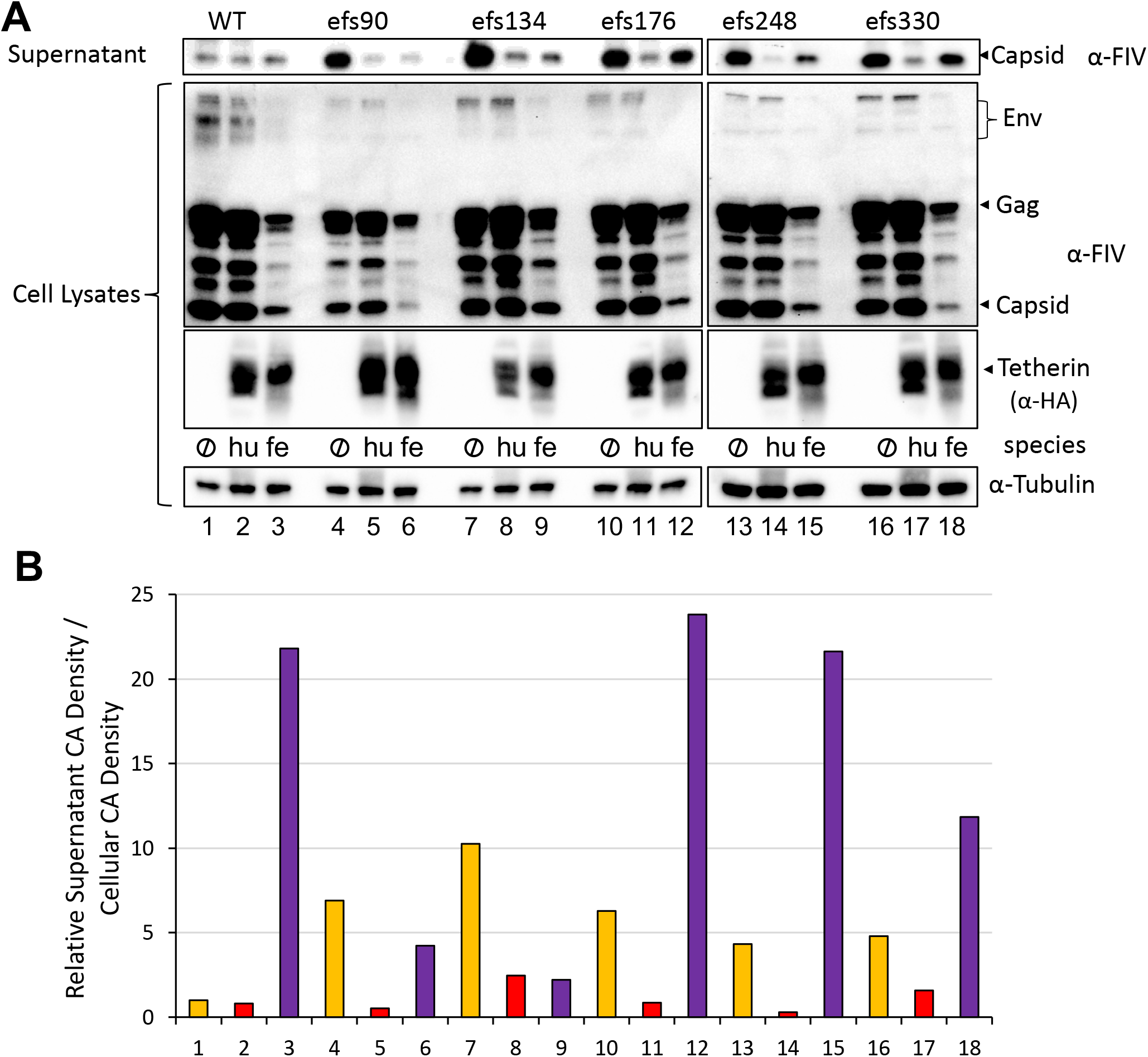
Effects of N-terminal deletions on feline tetherin restriction. **(A)** 293T cells having the indicated tetherin stably expressed under puromycin selection were transfected with the indicated pC36 proviral construct. Numbering indicates the last intact Env amino acid before early protein termination due to a frameshifting mutation. 48 hours after transfection cell lysates and supernatants were harvested and analyzed as in panel A. **(B)** Bands corresponding to FIV capsid from panel A were density quantified using ImageJ. The ratio of the supernatant to intracellular capsid band were calculated and normalized to FIVC36 from cells without tetherin. Lane numbers are the same as panel A. Experiment was repeated four times and a representative example is shown.

### Fess is necessary and sufficient to counteract tetherin

Considering these data, we hypothesized that the FIV Env signal sequence, which we designate Fess, is the necessary factor that mediates feline tetherin antagonism. We further hypothesized that it is sufficient (autonomously acting). Signal sequences, also known as leader sequences or signal peptides, are N-terminal peptides that were proposed in 1971 (Blobel and Sabatini, 1971) and subsequently established by Blöbel and colleagues (Blobel and Dobberstein, 1975a, b) to act as cellular “zip codes” that direct targeted translocation of newly synthesized proteins into the endoplasmic reticulum (ER) lumen in a signal recognition particle-dependent manner. Analogous subcellular targeting motifs, e.g., for mitochondria have also been described (Blobel, 2000). For enveloped viruses, signal sequences are the predominant mechanism targeting viral surface proteins to the correct cellular compartment to enable proper particle incorporation. In both eukaryotes and prokaryotes signal sequence lengths are generally very short, with a mean length of 23 +/− 6 amino acids (Hiss and Schneider, 2009)). Primate lentiviruses encode somewhat longer Env signal sequences, ranging between 19 and 45 amino acids in total length (**Figure S1**). Remarkably, our database searches and literature reviews indicate that FIV encodes the longest known signal sequence in any eukaryotic or prokaryotic species, or in any virus (175 amino acids). Despite the identification of this unusual property over 25 years ago (Pancino et al., 1993; Verschoor et al., 1993), the functional implications are unknown. To confirm that the signal sequence is directly responsible for enabling viral budding in the presence of domestic cat tetherin, and acts independently of other Env domains, we expressed it in trans, as a fusion to an irrelevant but trackable protein, GFP (**Figure 3A**). We co-expressed full-length Env, Fess-GFP, or a myc-epitope control protein (HIV Integrase) with FIVC36 efs90 (**Figure 3B**). Viral budding was assessed by immunoblotting supernatants with FIV antisera. As expected, FIVC36 efs90 budding was severely impaired by either human or feline tetherin (**Figure 3B**, top). In contrast, co-expression of either FIV Env or Fess-GFP rescued budding specifically in the presence of domestic cat tetherin but not human tetherin (**Figure 3B**, middle and bottom). These results highlight that Fess can function to counteract tetherin independently of other Env domains and is sufficient for FIV antagonism of tetherin.

**Figure 3.**
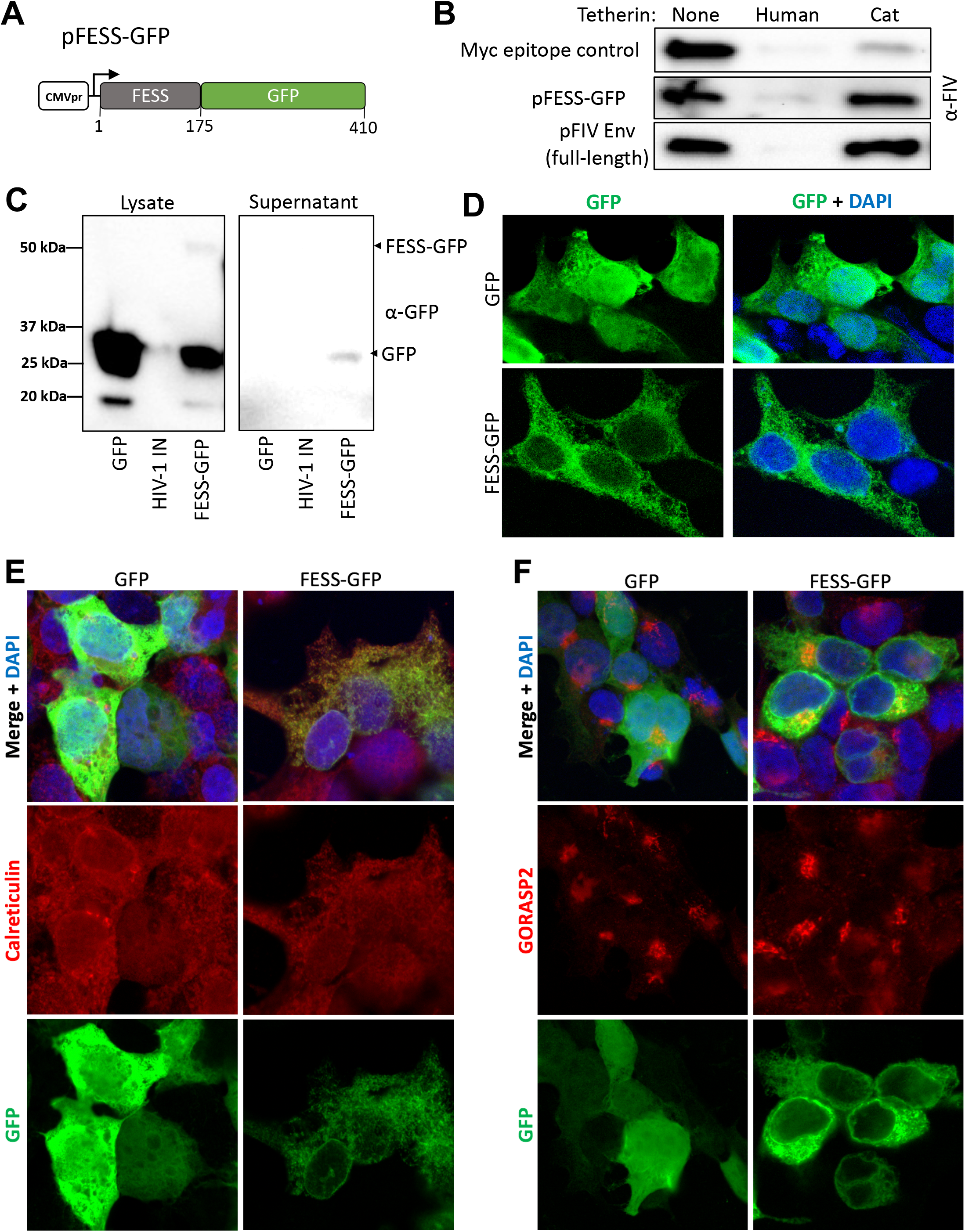
Fess is sufficient to counteract domestic cat tetherin and can direct protein translocation into the secretory pathway. **(A)** Diagram of the Fess-GFP construct. **(B)** 293T cells stably expressing tetherin were transfected as in Figure 1 with 1 μg pC36 EFS90 and 0.5 μg p1012-myc (control), p1012 SS-GFP, or pFE-C36 (full-length C36 Envelope). Equal volume supernatant was harvested 48 hours post-transfection, immunoblotted with cat sera reactive to FIV and viral capsid bands are shown. Experiment was repeated three times and a representative example is shown. 293T cells were transfected with plasmids that express eGFP, myc-tagged HIV-1 integrase (negative control for blot), or Fess-GFP (pEGFP-N1, p1012 -HIV-1-IN-myc, pFess-GFP) and cell lysates and supernatants were collected 48 hours post-transfection and immunoblotted with mouse anti-GFP and goat anti-mouse HRP. **(D-F)** Confocal microscopy of 293T cells transfected with pEGFP-N1 or pFess-GFP 16 hours post-transfection. Cells were mounted with ProLong Gold containing DAPI without additional staining **(D)** or were stained with rabbit anti-Calreticulin **(E)** or rabbit anti-GORASP2 **(F)** and Alexaflour 594 goat-anti-rabbit.

### Fess directs endoplasmic reticulum translocation and has dual function

To further characterize the Fess protein for localization properties and to test if it can fulfill traditional signal sequence functions in addition to acting as a tetherin antagonist, we also examined the fate of Fess-GFP. In immunoblots for GFP, although Fess-GFP was detectable, we observed intracellular accumulation of predominantly free GFP (**Figure 3C**, left), indicating that Fess is efficiently cleaved from GFP at the signal peptidase cleavage site. Free GFP was also detectable in the supernatant of Fess-GFP-transfected but not GFP-transfected cells, indicating Fess directed GFP translocation into the ER lumen and export via the secretory pathway (**Figure 3C**, right). Furthermore, immunofluorescence experiments were strongly suggestive of localization in the ER for the cleaved GFP signal in Fess-GFP expressing cells (**Figure 3D**). Confirming this, GFP co-localized with the ER-resident protein calreticulin (**Figure 3E**) but not the Golgi apparatus marker GORASP2 (**Figure 3F**). These experiments indicate that the majority of the free GFP seen in Figure 2C is located in the ER. Cumulatively, the data confirm that the FIV Env N-terminal 175 amino acids act as a bona fide signal sequence to direct protein translocation.

### Deletion of a central Fess motif has no effect on viral replication in the absence of tetherin but severely impairs virion release and viral replication in its presence

Although signal sequences show no conservation of amino acid sequences, they are functionally tripartite, with a positively charged N-terminal segment of variable length and sequence, a single central hydrophobic core (H region), and a short C-terminal and usually more conserved signal peptidase cleavage-site (Owji et al., 2018). In contrast to this typical architecture, we identified two distinct hydrophobic segments in Fess, which we designate H1 and H2 (**Figure 4A, B**). H1 is located in the central portion of Fess, downstream of Rev exon 1 and upstream of the cleavage site-adjacent H2 motif. To test the involvement of these regions in tetherin function, as well as viral viability and the processing and trafficking of Env, we generated in-frame deletions in the full-length virus. Mutant ΔH2/ΔC deletes the traditional signal sequence hydrophobic motif and adjacent cleavage signal. Mutant Δ40 was designed to delete much of the region (40 amino acids) between the end of Rev first exon residues and the onset of the traditional hydrophobic signal sequence and cleavage signal. The Δ40 deletion spans most of the H1 hydrophobic motif (VFS<u>ILYLFTGYIVYFL</u>) as well as a downstream region rich in charged (R, K, D, E) residues (**Figure 4A**). FIV C36Δ40 retained Env expression and normal replication kinetics in feline CrFK cells, whereas the FIV C36ΔH2/ΔC mutant had much lower Env protein expression in cells and greatly diminished replication (**Figure 4C, D**). Wild type FIV was inhibited by human but not feline tetherin as expected (**Figure 4E**). In contrast, the Δ40 mutation had two notable effects. It resulted in increased budding compared to wild type virus (compare black bars) in the absence of feline tetherin, but resulted in severe restriction in its presence (**Figure 4E**). Feline but not human tetherin restriction was selectively abrogated if these 40 amino acids were intact. These results, combined with the above observations, confirm that Fess confers resistance to domestic cat tetherin restriction of viral release from cells. Furthermore, the effects of Fess on Env SU/TM expression and trafficking to the cell surface are separable from its tetherin antagonism function.

**Figure 4.**
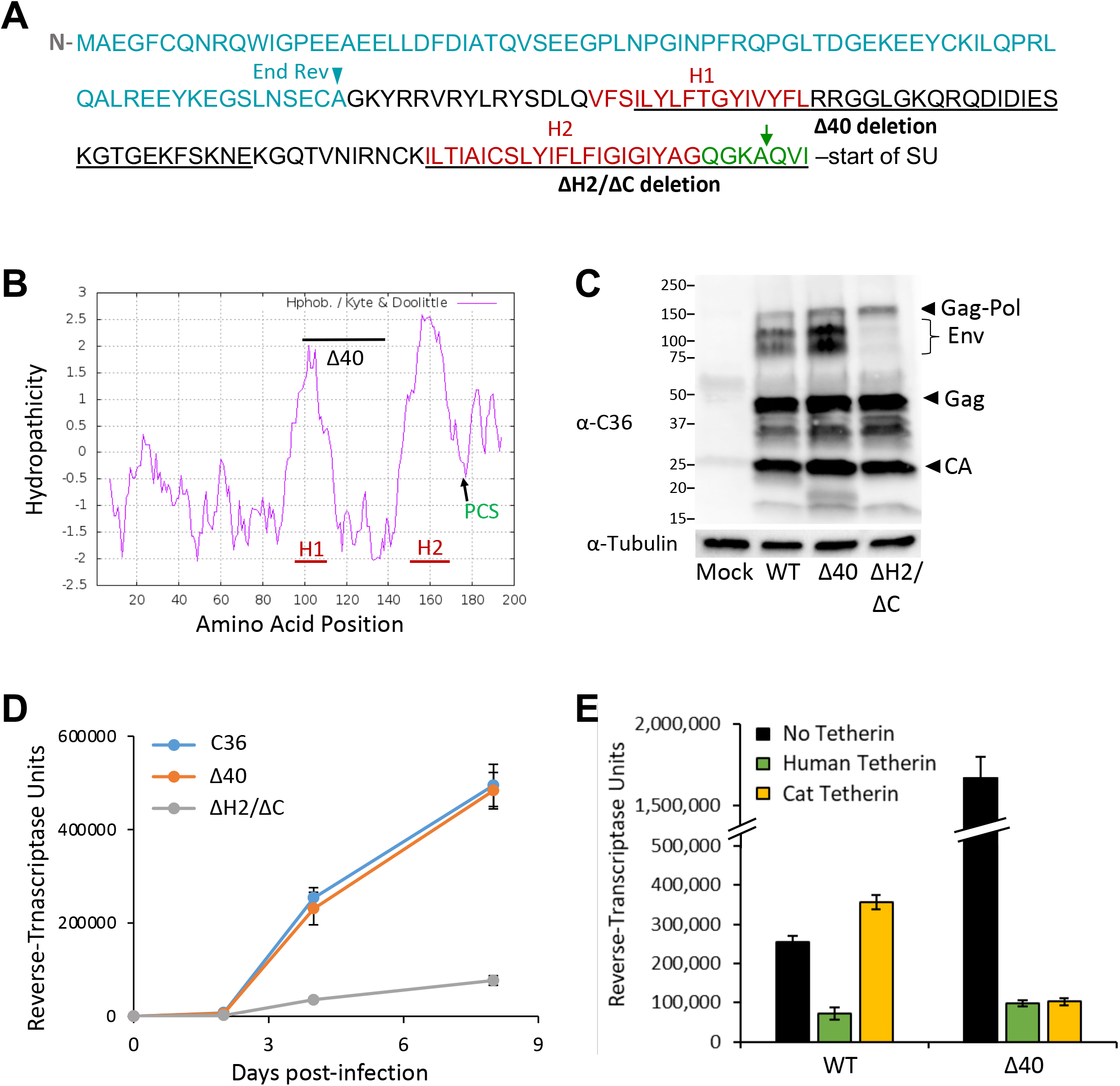
Fess has separable roles in Env expression and tetherin antagonism. **(A)** Fess amino acid sequence (FIV C36). Approximate hydrophobic regions H1 and H2 are indicated in red font, the C-region in green font, with a green arrow denoting the predicted signal peptidase cleavage site (Verschoor et al., 1993). The regions deleted in Δ40 and ΔH2/ΔC mutants of FIV C36 are underlined. Amino acids that are shared with Rev are in teal font. **(B)** FIV-C36 Fess amino acids 1-200 were plotted for hydropathicity using the Kyte and Doolittle model. H1: hydrophobic peak 1. H2: hydrophobic peak 2. PCS: predicted signal peptidase cleavage site. **(C)** 293T cells lacking tetherin were transfected with the indicated FIV-C36 proviral constructs, and 48 hours post-transfection cell lysates were harvested and immunoblotted with cat sera reactive to FIV. (C) CrFK cells stably expressing the FIV receptor CD134 were infected with the indicated viruses which were input normalized to reverse-transcriptase content. Cells were washed twice 16 hours post-infection and every other day 50 μL supernatant was collected for reverse-transcriptase quantification. Spreading replication was performed twice and one experiment is shown. **(E)** 293T cells with stable human or feline tetherin expression were transfected and analyzed as in Figure 2A. Reverse-transcriptase activity was quantified in each supernatant and the results are shown as means +/− standard deviations. This was repeated three times and a representative example is shown.

### Fess acts by blocking tetherin Incorporation into particles

Our previous data suggested that FIV antagonizes tetherin by a mechanism more similar to that of Ebola virus than that mediated by primate lentivirus accessory genes (Morrison et al., 2014). Both viruses act in a way that is linked to their respective Env glycoproteins but does not reduce cell surface or intracellular tetherin levels (Kuhl et al., 2011; Lopez et al., 2010; Morrison et al., 2014); see also **Figure 2A** here. This observation suggested to us a mechanism that acts locally, on a per-particle basis at the point of viral budding, to exclude the factor from the particle (Morrison et al., 2014). To determine whether Fess alters the particle association of tetherin, wild type or Δ40 FIV particles were produced in the presence or absence of stably expressed tetherins and then purified by ultracentrifugation over a sucrose cushion. Particles were then immunoblotted – using reverse transcriptase-normalized inputs – for tetherin. The results were dramatic. Wild-type FIVC36 virions contained minimal amounts of human or feline tetherin (**Figure 5**). In contrast, FIVC36Δ40 virions contained similarly low levels of human tetherin, but high levels of feline tetherin (**Figure 5**). Thus, the activity of Fess is both powerful and specific: intact Fess stringently blocks otherwise abundant virion incorporation of feline but not human tetherin. This mechanism is also unique among the lentiviral anti-tetherins.

**Figure 5.**
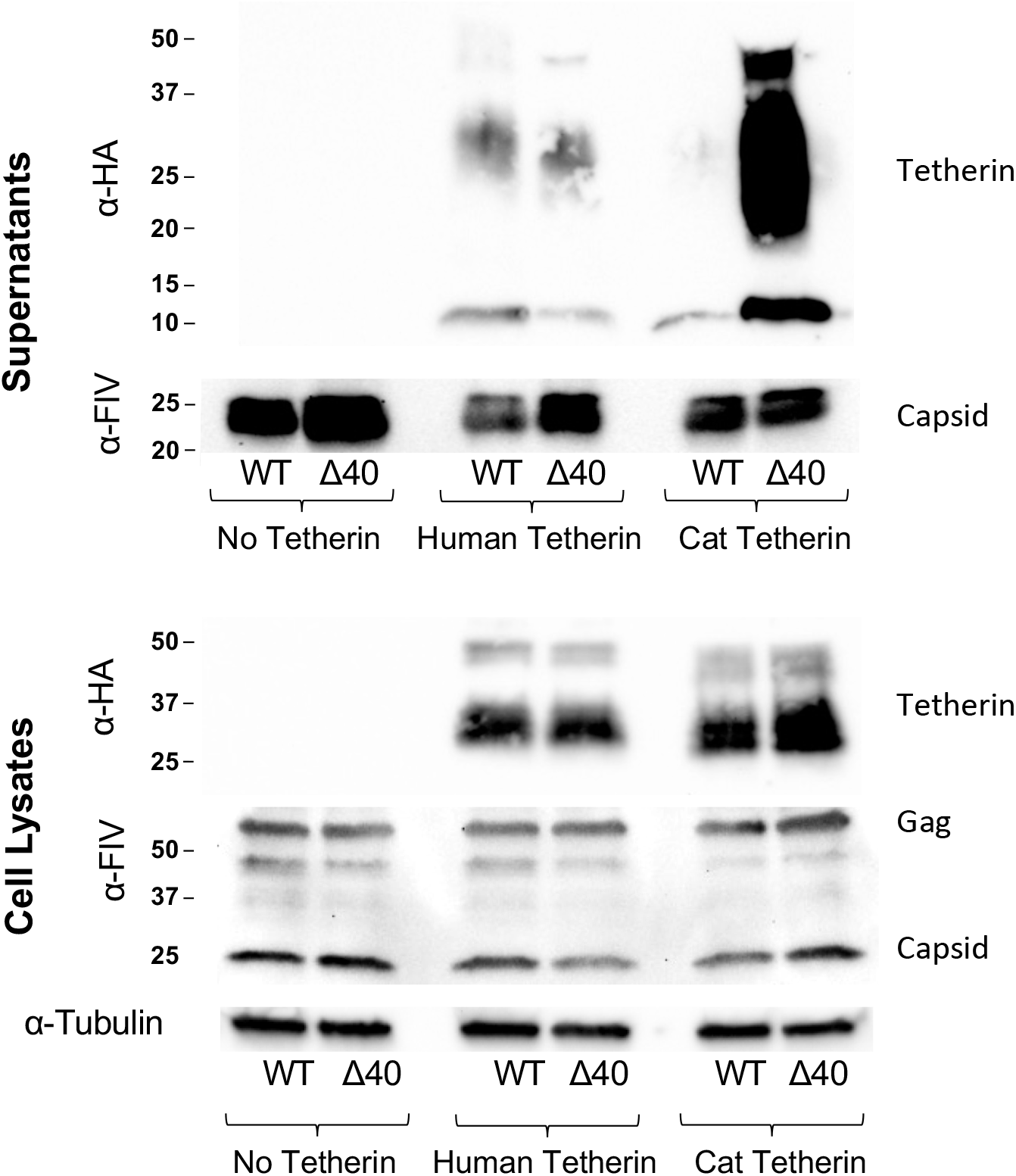
Fess blocks virion incorporation of feline tetherin. Control and HA-tetherin expressing 293T cells were transfected with indicated FIV proviral constructs as in Figure 2A. Supernatants were harvested and concentrated by ultracentrifugation over a 20% sucrose cushion. Concentrated virus was analyzed for reverse-transcriptase activity and RT-normalized inputs were used for immunoblotting with the indicated antibodies. A representative example is show for this experiment, which was repeated at least four times with similar results.

## DISCUSSION

Our results reveal that the Env glycoprotein signal sequence is the FIV tetherin antagonist, thus identifying the fourth lentiviral anti-tetherin protein and also, to the best of our knowledge, identifying the first new lentiviral accessory protein in decades. Almost all of Env – all of the SU and TM domains – was entirely dispensable for tetherin antagonism. Fess was both necessary and sufficient for enabling release of viral particles from tetherin expressing cells (**Figure 1–4**). We identified a central 40 amino acid segment of Fess that is located C-terminal to the first exon of Rev, is not needed for Env processing, and the function of which was previously unknown. Deletion of the segment did not affect FIV replication in the absence of tetherin but severely impaired budding of the virus in its presence. The mechanism fulfills multiple criteria for a specifically evolved restriction factor antagonism, as it is also virus- and species-specific: Fess did not protect FIV from human tetherin, or HIV-1 from either feline or human tetherin (**Figures 2 & 4E** and (Morrison et al., 2014)).

Simple retroviruses encode *gag*, *pol* and *env* genes that primarily encode the structural and enzymatic proteins needed for completing essential viral lifecycle steps. Lentiviruses, by contrast, are complex retroviruses that establish persistent, lifelong infections and must therefore evade innate and adaptive immunity for years. To meet this challenge, they have evolved additional accessory proteins that modulate host immune responses and antagonize innate immune effectors. HIV-1 and simian lentiviruses variably use the small accessory proteins Vpu and Nef to overcome the block to nascent viral budding imposed by tetherin, which appears to have afforded genetic flexibility. The evolution of these proteins not only underlies persistence in a given species, but has also been shown to be critical for host switching, as in the case of the acquisition of Vpu during the adaptation of SIVcpz to become HIV-1 (Neil, 2017; Sauter et al., 2009). Presumably due to the pressures posed by such viral proteins and also from non-retroviral antagonists, tetherin proteins show evidence of positive selection in mammals (Lim et al., 2010; Liu et al., 2010; McNatt et al., 2009).

Non-primate lentiviruses encode more limited accessory gene repertoires than primate lentiviruses and they specifically lack Vpu and Nef proteins. The prior mapping of anti-tetherin activity to the *Env* gene of FIV suggested antagonism by the full Env glycoprotein similar to what has been observed for HIV-2 (Celestino et al., 2012; Dietrich et al., 2011; Le Tortorec and Neil, 2009; Morrison et al., 2014). However, our data reveal that FIV SU and TM are dispensable. We showed previously that FIV does not degrade tetherin or down-regulate it from the cell surface (Morrison et al., 2014), identifying a contrast with primate lentiviral Vpu, Nef or Env proteins, which all mediate functional depletion of tetherin from the primary site of viral budding via intracellular sequestration, endocytosis and lysosomal degradation of the protein (Jia et al., 2009; Le Tortorec and Neil, 2009; Zhang et al., 2009). Here we show that Fess instead excludes tetherin from the particle, establishing a mechanism unique among the lentiviruses (**Figure 5**). Fess possesses autonomous restriction blocking activity and can also direct the ER translocation and export of an unrelated protein (**Figure 3**).

Signal sequences (signal peptides) act as intracellular zip codes that direct the location of many cellular proteins that are destined for extra-cytoplasmic locations. Proteins destined for secretion or plasma membrane residence are translocated as preproteins through or into the ER membrane. After preprotein translocation has partially completed, the signal peptidase enzyme cleaves the signal sequence away, which enables correct folding of the mature protein. Signal sequences are generally quite short (under 30 amino acids), do not have further functions, and are mostly degraded in short order by the signal peptide peptidase. In the case of RNA viruses, however, genome sizes are severely constrained and, for retroviruses, genetic efficiency in the form of overlapping reading frames and multiple purpose proteins, e.g., Nef, are observed. Dual purposing of the signal sequence by FIV is an interesting example. The amino acids encoded by the first exon of Rev are also present in the first 80 amino acids of Fess, which adds a third function to this compressed region of the genome. There are a few prior examples of cellular protein signal sequences with additional cellular functions (Martoglio and Dobberstein, 1998) and a few in other viruses as well. However, the latter generally involve direct participation in mechanics of the virus’s replication machinery. Following signal peptidase processing of the Arenavirus envelope glycoprotein, the signal sequence is not degraded and instead forms a tripartite complex with the mature glycoprotein subunits, which is necessary for glycoprotein mediated fusion with target cells (Nunberg and York, 2012). In this case the signal sequence function remains tied to that of the envelope protein. Among retroviruses, the spumaretrovirus foamy virus signal sequence binds to cognate Gag molecules and is packaged into viral particles, where it appears to be necessary for proper virion morphogenesis (Geiselhart et al., 2003; Lindemann et al., 2001). The signal sequences of the betaretroviruses mouse mammary tumor virus (Dultz et al., 2008) and Jaagsiekte sheep retrovirus (Caporale et al., 2009) Env proteins traffic to the nucleoli of infected cells, where they are involved in modulating nuclear export of unspliced viral mRNAs.

In the case of Fess, the signal sequence has evolved to counter a main host defense and can properly be considered a viral accessory protein. We propose that Env signal sequences with additional functions independent of directing Env translocation may in fact be a more general non-primate lentivirus property, since the signal sequences of these viruses vary in length but are all significantly longer (66-176 amino acids, **Figure S1**) than the signal sequences of primate lentiviruses and most cellular signal sequences (Martoglio and Dobberstein, 1998; Pancino et al., 1994). Indeed, the full length equine infectious anemia virus Env glycoprotein has been reported to counteract horse tetherin, also by a mechanism that does not degrade the factor; whether all or just part of Env is the antagonist has not been determined (Yin et al., 2014).

Our results underscore the centrality of tetherin to mammalian defense against lentiviruses in widely different circumstances. Feline and primate lentiviruses share distant ancestry, with the former likely to have colonized some but not all feline lineages sometime after the modern felid species radiation in the late Miocene, c.a. 11 million years ago (Mya) (Pecon-Slattery et al., 2008). Major commonalities do persist between FIV and HIV in pathophysiology (AIDS) and dependency factor utilization, such as CXCR4 and LEDGF, and both have Vif proteins that degrade APOBEC3 proteins (Llano et al., 2006; Münk et al., 2008; Poeschla and Looney, 1998). Domestic cat FIV is an AIDS-causing lentivirus like HIV-1, yet it and its ancestral felid species relatives have been on an independent evolutionary trajectory for millions of years. On the host side of the equation, the tiger and the domestic cat tetherin proteins share a likely more ancient (c.a. 60 to 11 Mya) truncation of the cytoplasmic tail, with the loss of 19 of 27 amino acids, including a dual tyrosine motif (Morrison et al., 2014). The parallel evolution by FIV of an anti-tetherin protein that is structurally, functionally and mechanistically very different from those of the primate lentivirus proteins in consistent with this and other evidence for the genetic plasticity of this host factor. The unusual architecture of tetherins rather than primary sequence is critical for their function, as well as their versatility against other groups of enveloped viruses (Blanco-Melo et al., 2016; Heusinger et al., 2015; Perez-Caballero et al., 2009). Investigation of other lentiviruses may uncover further viral solutions to the problem of tetherin.

## MATERIALS AND METHODS

### Cells

239T and Crandell feline kidney (CrFK) cells were cultured in DMEM with 10% fetal calf serum (FBS), penicillin-streptomycin, and L-glutamine. Stable HA-tetherin expressing cells have been previously described (Morrison et al., 2014) and were additionally cultured in 3 μg/mL puromycin.

### Vectors, viruses and plasmids

pCT-C36^A+^ and pFE-C36 (subclone encoding Env protein) were used as the basis for mutagenesis (Morrison et al., 2014). They employ the 5 ‘U3-replacement strategy that enabled FIV production in human cells, in which the FIV U3 has virtually no promoter function) (Poeschla and Looney, 1998; Poeschla et al., 1998). In this case we applied this to the proviral clone C36 (de Rozieres et al., 2004), Additionally C36 *Env* was subcloned from NheI-digested pCT-C36A+ into the NheI site of gammaretroviral vector pJZ308 (Poeschla and Looney, 1998) to yield pJZC36. Env-frameshift mutants of C36 were constructed within pFE-C36 by site directed mutagenesis (Efs330) or overlap-extension PCR between NotI and MfeI restriction enzyme sites, approximately comprising *Env* amino acids 1-500. Insertions to frameshift Env are denoted here (lower-case letter indicates inserted nucleotide, enzyme in parentheses indicates restriction site added by nucleotide insertion):

Efs90: GGTAAGATATTTAAGATAtCTCTGATTTACAAGTATTTAG (EcoRV)
Efs134: CTGGGGAAAAATTTAatTAAAAATGAAAAGGGAC (PacI)
Efs176: GACAAGGTAAGGCACAAGctTAATATGGAGACTCCCACCC (HindIII)
Efs247: GAAAGCTACAAGAtAATcTAGAAGGGGAAAAGTTTGG (XbaI)
Efs330: CAAATCCCACTGATCAATTAgtcgacagTACATTTGGACCTAATC (ScaI)

Mutants were verified by restriction enzyme digestion and sequencing across the cloned fragment. An AvrII/BglII fragment from pFE-C36 was then moved into pJZ C36 (AvrII/BglII), then an NheI fragment from pJZ C36 was then swapped into an NheI digested pCT-C36 to create pCT-C36 mutants with the indicated insertions causing *Env* frame-shift but no other changes. Each mutant was again verified by sequencing across the entire NheI fragment. C36Δ40 and C36ΔH2/ΔC were cloned by overlap extension PCR to remove 40 amino acids (residues 99-138) or 29 amino acids (ΔH2/ΔC, residues 150-178), and re-inserted between AvrII and EcoNI digested pJZ C36, then an NheI fragment of pJZC36Δ40 or ΔH2 containing C36 *Env* was again inserted back into NheI digested pCT-C36^A+^. Codon optimized Fess (amino acids 1-178) was synthesized as a gBlock (IDT). Complementary Fess and GFP cDNAs were generated by PCR and cloned into a NotI/BglII digested p1012 IN-myc (Vanegas et al., 2005) using the GeneArt Seamless Cloning System (ThermoFisher), with a single amino acid linker (S) separating Fess and GFP.

### Transfections and particle analyses

4×10^5^ 293T cells were plated per well of a 6-well plate, allowed to adhere overnight, and PEI transfected. 1.5 g total DNA was added to 35 μL of Optimem without serum and 6 μL of 1 μg/μL PEI before brief vortexing and incubation at room temperature for 30min. Transfection mix was added drop-wise to cells and washed with fresh complete media after 8-16 hours. 48 hours post-transfection, supernatant was harvested and filtered through a 0.45 μM filter. For analysis of tetherin incorporation into viral particles, supernatant was concentrated by ultracentrifugation over a 20% sucrose cushion. At the time of supernatant harvest, cells were lysed in 1x radioimmunoprecipitation (RIPA) buffer (150 mM NaCl, 0.5% deoxycholate, 0.1% sodium dodecyl sulfate, 1% NP-40, 150 mM Tris-HCl pH8.0). Immunoblotting was performed with cat serum reactive to FIV-PPR (gift of Peggy Barr), rat α-HA (Roche), mouse anti-GFP (JL-8 clone, Takara Bio) or mouse anti-alpha-tubulin (Sigma). Immunoblot band density was quantified using ImageJ.

### Reverse-transcriptase activity

Reverse-transcriptase (RT) activity was quantified by use of a real-time PCR assay as previously described (Vermeire et al., 2012). 5 μL of supernatant was mixed with 5 μL of 2x viral lysis buffer (0.25% Triton X-100, 50 mM KCL, 100 mM TrisHCL, pH 7.4, 40% glycerol, and 2% v/v RNAse inhibitor) and incubated at room temperature for 10 minutes, then 90 μL of sterile water was added. Samples were diluted 1:100 in sterile water, then 9 μL of diluted, lysed sample was used in a 20 μL qPCR reaction containing 10 μL of 2x SYBR Green master mix (Apex Sybr Green, Quintarabio), 120 nM MS2 cDNA primers, and 0.055 A_260_ units of MS2 RNA (Sigma, catalog # 10165948001).

### Immunofluorescence

1 x 10^5^ 293T cells were plated on LabTek II chamber slides, allowed to adhere overnight and transfected with 500 ng indicated plasmids (pEGFP-N1 or pFess-GFP). 48 hours post-transfection cells were fixed with 4% (wt/vol) paraformaldehyde for 10 minutes at room temperature, permeabilized with methanol, stained for 1 hour at room temperature with rabbit anti-Calreticulin (1:50, Abcam ab2907) or anti-GORASP2 (1:100, Sigma HPA035274), washed 3x with PBS and stained for 1 hour at room temperature with Alexafluor 594-anti-rabbit-IgG (1:500). Wells were again washed 3x with PBS then mounted with ProLong Gold antifade reagent with DAPI (Invitrogen P36935). Images were collected on a Zeiss LSM780.

## ACKNOWLEDGMENTS

Supported by NIH grant AI77344 and DP1DA043915. Imaging experiments used the Advanced Light Microscopy Core at the Anschutz Medical Campus, which is supported by NIH NS048154 and DK116073. We thank other laboratory members for helpful suggestions.

## AUTHOR CONTRIBUTIONS

J.H.M and E.M.P formulated ideas, hypotheses, and experimental approaches. J.H.M performed experiments. J.H.M and E.M.P analyzed and interpreted data and wrote the manuscript.

---

**Figure S1.**
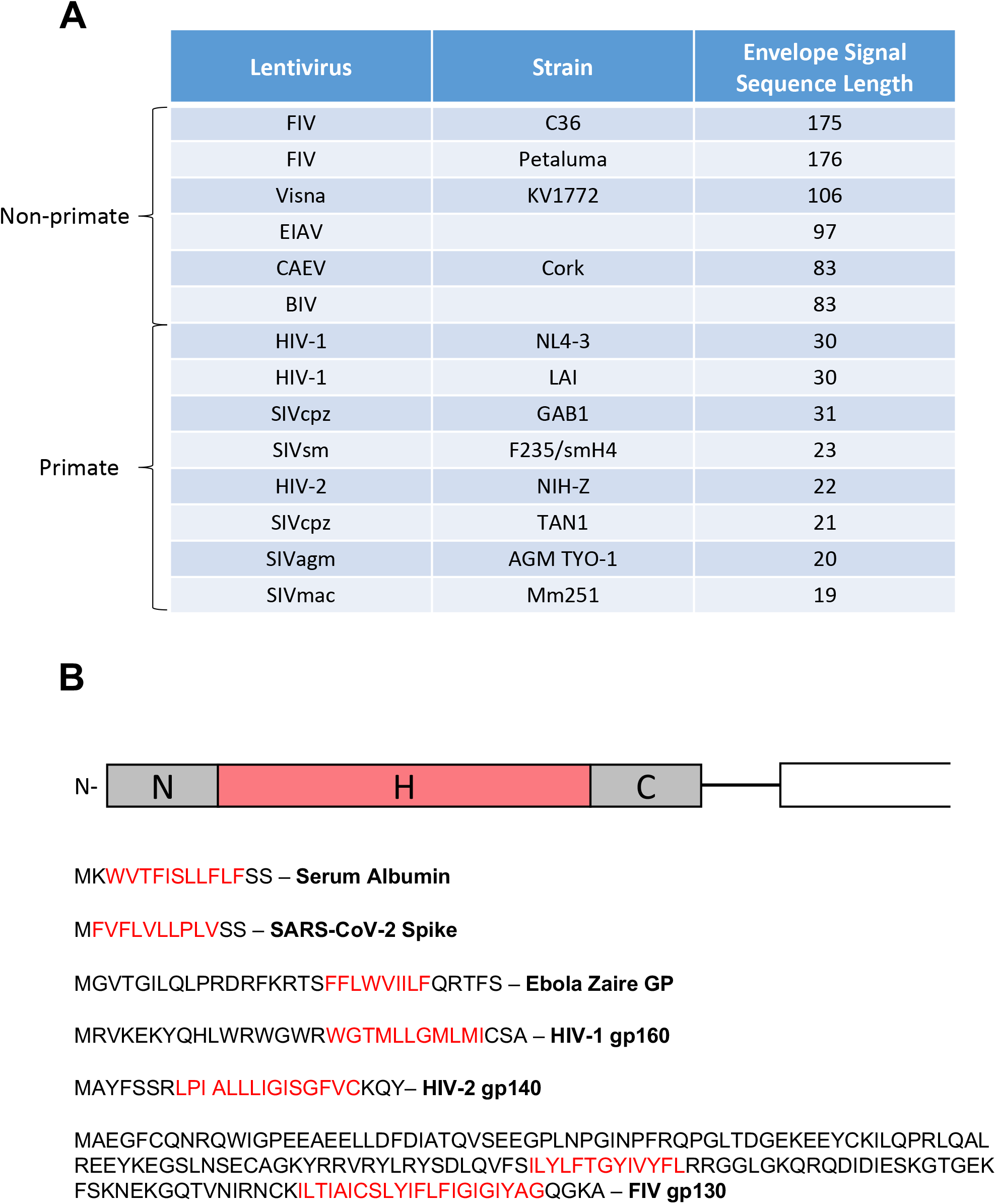
Signal sequence lengths. Signal sequence lengths reported are from manual assertion according to sequence analysis in the Uniprot database (UniProt Consortium, 2018) or were previously analyzed using the PCgene program (Pancino et al., 1994). **(A)** Lentiviruses. **(B)** Human albumin, SARS-CoV-2 Spike, Ebola GP, HIV-2 Env, HIV-1 Env, and FIV Env signal peptides. Hydrophobic regions are shown in red font.

## REFERENCES

Blanco-Melo, D., Venkatesh, S., and Bieniasz, P.D. (2016). Origins and Evolution of tetherin, an Orphan Antiviral Gene. Cell host & microbe 20, 189–201.

Blobel, G. (2000). Protein targeting (Nobel lecture). Chembiochem 1, 86–102.

Blobel, G., and Dobberstein, B. (1975a). Transfer of proteins across membranes. I. Presence of proteolytically processed and unprocessed nascent immunoglobulin light chains on membrane-bound ribosomes of murine myeloma. The Journal of cell biology 67, 835–851.

Blobel, G., and Dobberstein, B. (1975b). Transfer of proteins across membranes. II. Reconstitution of functional rough microsomes from heterologous components. The Journal of cell biology 67, 852–862.

Blobel, G., and Sabatini, D. (1971). Ribosome-membrane interaction in eukaryotic cells. In Biomembranes, L. Manson, ed. (Plenum Publishing Corporation), pp. 193–195.

Caporale, M., Arnaud, F., Mura, M., Golder, M., Murgia, C., and Palmarini, M. (2009). The signal peptide of a simple retrovirus envelope functions as a posttranscriptional regulator of viral gene expression. Journal of virology 83, 4591–4604.

Celestino, M., Calistri, A., Del Vecchio, C., Salata, C., Chiuppesi, F., Pistello, M., Borsetti, A., Palu, G., and Parolin, C. (2012). Feline tetherin is characterized by a short N-terminal region and is counteracted by the feline immunodeficiency virus envelope glycoprotein. Journal of virology 86, 6688–6700.

Cocka, L.J., and Bates, P. (2012). Identification of alternatively translated Tetherin isoforms with differing antiviral and signaling activities. PLoS pathogens 8, e1002931.

de Rozieres, S., Mathiason, C.K., Rolston, M.R., Chatterji, U., Hoover, E.A., and Elder, J.H. (2004). Characterization of a highly pathogenic molecular clone of feline immunodeficiency virus clade C. Journal of virology 78, 8971–8982.

Dietrich, I., McMonagle, E.L., Petit, S.J., Vijayakrishnan, S., Logan, N., Chan, C.N., Towers, G.J., Hosie, M.J., and Willett, B.J. (2011). Feline tetherin efficiently restricts release of feline immunodeficiency virus but not spreading of infection. Journal of virology 85, 5840–5852.

Dultz, E., Hildenbeutel, M., Martoglio, B., Hochman, J., Dobberstein, B., and Kapp, K. (2008). The signal peptide of the mouse mammary tumor virus Rem protein is released from the endoplasmic reticulum membrane and accumulates in nucleoli. The Journal of biological chemistry 283, 9966–9976.

Galao, R.P., Le Tortorec, A., Pickering, S., Kueck, T., and Neil, S.J. (2012). Innate sensing of HIV-1 assembly by Tetherin induces NFkappaB-dependent proinflammatory responses. Cell host & microbe 12, 633–644.

Geiselhart, V., Schwantes, A., Bastone, P., Frech, M., and Lochelt, M. (2003). Features of the Env leader protein and the N-terminal Gag domain of feline foamy virus important for virus morphogenesis. Virology 310, 235–244.

Heusinger, E., Kluge, S.F., Kirchhoff, F., and Sauter, D. (2015). Early Vertebrate Evolution of the Host Restriction Factor Tetherin. Journal of virology 89, 12154–12165.

Hirsch, V.M., Olmsted, R.A., Murphey-Corb, M., Purcell, R.H., and Johnson, P.R. (1989). An African primate lentivirus (SIVsm) closely related to HIV-2. Nature 339, 389–392.

Hiss, J.A., and Schneider, G. (2009). Domain organization of long autotransporter signal sequences. Bioinform Biol Insights 3, 189–204.

Jia, B., Serra-Moreno, R., Neidermyer, W., Rahmberg, A., Mackey, J., Fofana, I.B., Johnson, W.E., Westmoreland, S., and Evans, D.T. (2009). Species-specific activity of SIV Nef and HIV-1 Vpu in overcoming restriction by tetherin/BST2. PLoS pathogens 5, e1000429.

Kuhl, A., Banning, C., Marzi, A., Votteler, J., Steffen, I., Bertram, S., Glowacka, I., Konrad, A., Sturzl, M., Guo, J.T., et al. (2011). The Ebola virus glycoprotein and HIV-1 Vpu employ different strategies to counteract the antiviral factor tetherin. J Infect Dis 204 Suppl 3, S850–860.

Le Tortorec, A., and Neil, S.J. (2009). Antagonism to and intracellular sequestration of human tetherin by the human immunodeficiency virus type 2 envelope glycoprotein. Journal of virology 83, 11966–11978.

Lim, E.S., Malik, H.S., and Emerman, M. (2010). Ancient adaptive evolution of tetherin shaped the functions of Vpu and Nef in human immunodeficiency virus and primate lentiviruses. J Virol 84, 7124–7134.

Lindemann, D., Pietschmann, T., Picard-Maureau, M., Berg, A., Heinkelein, M., Thurow, J., Knaus, P., Zentgraf, H., and Rethwilm, A. (2001). A particle-associated glycoprotein signal peptide essential for virus maturation and infectivity. Journal of virology 75, 5762–5771.

Liu, J., Chen, K., Wang, J.H., and Zhang, C. (2010). Molecular evolution of the primate antiviral restriction factor tetherin. PloS one 5, e11904.

Llano, M., Saenz, D.T., Meehan, A., Wongthida, P., Peretz, M., Walker, W.H., Teo, W., and Poeschla, E.M. (2006). An Essential Role for LEDGF/p75 in HIV Integration. Science 314, 461–464.

Lopez, L.A., Yang, S.J., Hauser, H., Exline, C.M., Haworth, K.G., Oldenburg, J., and Cannon, P.M. (2010). Ebola virus glycoprotein counteracts BST-2/Tetherin restriction in a sequence-independent manner that does not require tetherin surface removal. Journal of virology 84, 7243–7255.

Martoglio, B., and Dobberstein, B. (1998). Signal sequences: more than just greasy peptides. Trends Cell Biol 8, 410–415.

McNatt, M.W., Zang, T., Hatziioannou, T., Bartlett, M., Fofana, I.B., Johnson, W.E., Neil, S.J., and Bieniasz, P.D. (2009). Species-specific activity of HIV-1 Vpu and positive selection of tetherin transmembrane domain variants. PLoS Pathog 5, e1000300.

Miyakawa, K., Ryo, A., Murakami, T., Ohba, K., Yamaoka, S., Fukuda, M., Guatelli, J., and Yamamoto, N. (2009). BCA2/Rabring7 promotes tetherin-dependent HIV-1 restriction. PLoS pathogens 5, e1000700.

Morrison, J.H., Guevara, R.B., Marcano, A.C., Saenz, D.T., Fadel, H.J., Rogstad, D.K., and Poeschla, E.M. (2014). Feline immunodeficiency virus envelope glycoproteins antagonize tetherin through a distinctive mechanism that requires virion incorporation. Journal of virology 88, 3255–3272.

Münk, C., Beck, T., Zielonka, J., Hotz-Wagenblatt, A., Chareza, S., Battenberg, M., Thielebein, J., Cichutek, K., Bravo, I.G., O’Brien, S.J., et al. (2008). Functions, structure, and read-through alternative splicing of feline APOBEC3 genes. Genome Biol 9, R48.

Neil, S.J. (2017). Exercising Restraint. Cell Host Microbe 21, 274–277.

Neil, S.J., Zang, T., and Bieniasz, P.D. (2008). Tetherin inhibits retrovirus release and is antagonized by HIV-1 Vpu. Nature 451, 425–430.

Nunberg, J.H., and York, J. (2012). The curious case of arenavirus entry, and its inhibition. Viruses 4, 83–101.

Owji, H., Nezafat, N., Negahdaripour, M., Hajiebrahimi, A., and Ghasemi, Y. (2018). A comprehensive review of signal peptides: Structure, roles, and applications. Eur J Cell Biol 97, 422–441.

Pancino, G., Ellerbrok, H., Sitbon, M., and Sonigo, P. (1994). Conserved framework of envelope glycoproteins among lentiviruses. Curr Top Microbiol Immunol 188, 77–105.

Pancino, G., Fossati, I., Chappey, C., Castelot, S., Hurtrel, B., Moraillon, A., Klatzmann, D., and Sonigo, P. (1993). Structure and variations of feline immunodeficiency virus envelope glycoproteins. Virology 192, 659–662.

Pecon-Slattery, J., Troyer, J.L., Johnson, W.E., and O’Brien, S.J. (2008). Evolution of feline immunodeficiency virus in Felidae: implications for human health and wildlife ecology. Veterinary immunology and immunopathology 123, 32–44.

Perez-Caballero, D., Zang, T., Ebrahimi, A., McNatt, M.W., Gregory, D.A., Johnson, M.C., and Bieniasz, P.D. (2009). Tetherin inhibits HIV-1 release by directly tethering virions to cells. Cell 139, 499–511.

Poeschla, E., and Looney, D. (1998). CXCR4 is required by a non-primate lentivirus: heterologous expression of feline immunodeficiency virus in human, rodent and feline cells. Journal of virology 72, 6858–6866.

Poeschla, E., Wong-Staal, F., and Looney, D. (1998). Efficient transduction of nondividing cells by feline immunodeficiency virus lentiviral vectors. Nature medicine 4, 354–357.

Sauter, D., Schindler, M., Specht, A., Landford, W.N., Munch, J., Kim, K.A., Votteler, J., Schubert, U., Bibollet-Ruche, F., Keele, B.F., et al. (2009). Tetherin-driven adaptation of Vpu and Nef function and the evolution of pandemic and nonpandemic HIV-1 strains. Cell host & microbe 6, 409–421.

UniProt Consortium, T. (2018). UniProt: the universal protein knowledgebase. Nucleic acids research 46, 2699.

Van Damme, N., Goff, D., Katsura, C., Jorgenson, R.L., Mitchell, R., Johnson, M.C., Stephens, E.B., and Guatelli, J. (2008). The interferon-induced protein BST-2 restricts HIV-1 release and is downregulated from the cell surface by the viral Vpu protein. Cell host & microbe 3, 245–252.

Vanegas, M., Llano, M., Delgado, S., Thompson, D., Peretz, M., and Poeschla, E. (2005). Identification of the LEDGF/p75 HIV-1 integrase-interaction domain and NLS reveals NLS-independent chromatin tethering. J Cell Sci 118, 1733–1743.

Vermeire, J., Naessens, E., Vanderstraeten, H., Landi, A., Iannucci, V., Van Nuffel, A., Taghon, T., Pizzato, M., and Verhasselt, B. (2012). Quantification of reverse transcriptase activity by real-time PCR as a fast and accurate method for titration of HIV, lenti- and retroviral vectors. PloS one 7, e50859.

Verschoor, E.J., Hulskotte, E.G., Ederveen, J., Koolen, M.J., Horzinek, M.C., and Rottier, P.J. (1993). Post-translational processing of the feline immunodeficiency virus envelope precursor protein. Virology 193, 433–438.

Yin, X., Hu, Z., Gu, Q., Wu, X., Zheng, Y.H., Wei, P., and Wang, X. (2014). Equine tetherin blocks retrovirus release and its activity is antagonized by equine infectious anemia virus envelope protein. J Virol 88, 1259–1270.

Zhang, F., Wilson, S.J., Landford, W.C., Virgen, B., Gregory, D., Johnson, M.C., Munch, J., Kirchhoff, F., Bieniasz, P.D., and Hatziioannou, T. (2009). Nef proteins from simian immunodeficiency viruses are tetherin antagonists. Cell host & microbe 6, 54–67.

